# Population genomic analysis reveals high inbreeding in a Hawksbill turtle population nesting in Singapore

**DOI:** 10.1101/2023.12.28.573525

**Authors:** Regine Hui Yi Tiong, Justine Dacanay, Akira Uchida, Archita Sanjay Desai, Namrata Kalsi, Collin Tong, Hie Lim Kim

**Affiliations:** Singapore Centre for Environmental Life Sciences Engineering, Nanyang Technological University, Singapore; Asian School of the Environment, Nanyang Technological University, Singapore; Coastal & Marine Branch, National Parks Board, Singapore

## Abstract

Seven species of marine turtles remain in a world currently threatened by anthropogenic activities and climate change, standing at the precipice of extinction. Urgent conservation endeavours are imperative to safeguard their survival and preserve the biodiversity of marine ecosystems. Yet, genetic studies on these turtles have leaned on restricted genetic markers, such as the mitochondrial control region. The markers could provide incomplete or biased estimations of genetic diversity and population structure, thereby limiting precise conservation strategies. Here, we have generated a *de novo* genome assembly and high-quality whole-genome population datasets from hawksbill turtles nesting and foraging in Singapore. This initiative aims to contribute to unbiased, fine-resolution genetic data on the species, conducting a comprehensive population genomic study. Our analysis results demonstrated a remarkable enhancement in genetic markers. While we identified five different haplotypes defined by five variants within 69 mitochondrial control region sequences, the analysis of 35 whole genome sequences uncovered approximately 12 million single nucleotide polymorphisms (SNPs). Within the Singapore hawksbill turtle population, our whole-genome analysis revealed a pronounced degree of inbreeding, with most samples sharing at least a first cousin relationship. Furthermore, within a multiple-paternity nest, we identified related parents. Additionally, our inference of demographic history underscored the impact of past climate change on the decreasing hawksbill turtle population. We believe this pioneering study will substantially enhance the field of conservation genetic study of marine turtles.

## Introduction

Marine turtles are ancient species which have lived since the era of dinosaurs 200 million years ago, as evidenced by paleontological records such as the Archelon (Wieland, 1896). Despite their historical resilience, marine turtles have been facing risk of extinction. Only seven extant species remain, and all are endangered. The decline of marine turtle populations, leading to the potential extinction, resonates with the overwhelming impact of anthropogenic activities on their habitat (McClenachan et al., 2006), overexploitation for their meat and intricately patterned carapace (Parsons, 1972), and the ornaments crafted from their shells (Meylan & Donnelly, 1999). However, the current greatest threat to marine turtle populations is climate change. Their unique life cycle renders them vulnerable to environmental change. Rising temperatures skew the sex ratios of turtle populations due to temperature-dependent sex determination during the development of turtle embryos (Bull & Vogt, 1979). Sea level rise inundates nests, reducing egg viability (Pike et al., 2015) and eventually lead to the loss of nesting habitats (Fish et al., 2005).

Among the seven marine turtle species, the hawksbill turtle stands at the most critically endangered status (Meylan & Donnelly, 1999). The species is distributed in the Indo-Pacific and Atlantic regions (Arantes et al., 2020) which is crucial for biodiversity hotspots. In these regions, hawksbill turtles occupy a pivotal ecological position in both marine and coastal ecosystems. In marine environments, they play a significant role in preserving coral reef ecosystems. By selectively feeding on sponges, the turtles mitigate the proliferation of sponges (León & Bjorndal, 2002) that compete with corals for space, thereby safeguarding marine life relying on coral reefs (Hill, 1998). The declining population of hawksbill turtles has led to the loss of coral reef ecosystems (Jackson, 1997). In coastal ecosystems, hawksbill turtle eggs serve as rich nutrients in impoverished sandy beach environments, contributing to the intricate web of coastal life (Leighton et al., 2011). The intricate and irreplaceable role of marine hawksbill turtles in ecosystems underscores the urgency of aiding the recovery of this critically endangered species to conserve marine biodiversity in Indo-Pacific regions.

Understanding the genetics of hawksbill turtles will unravel their evolutionary history over millennia and shed light on the current status of the population in the face of climate change. Genetics has emerged as a powerful tool in shaping effective conservation strategies (Steiner et al., 2013) by providing knowledge of the level of genetic diversity and population structure (Hedrick, 2001). Traditionally, conservation genetic studies have relied on genetic markers such as mitochondrial control regions or nuclear microsatellites (Jensen et al., 2013). A recently announced project aimed at constructing a database of hawksbill mitochondrial DNA sequences was initiated to track illegally traded hawksbill shells (LaCasella et al., 2021).

However, those genetic markers could result to estimation of an incomplete or biased population history, potentially limiting the precision of conservation strategies. Whole genome sequencing data is high-resolution genetic markers, enabling an unbiased and comprehensive assessment of genetic diversity - a crucial approach for conservation genetic studies. While the first *de novo* genome assemblies of two marine turtle species, the green and leatherback turtle, were published (Bentley et al., 2022), their genome analysis revealed an unexpectedly small genetic diversity in leatherback turtles compared to green turtles. Recently, a *de novo* genome assembly of the hawksbill turtle was reported, and comparative genome analysis with other marine turtle species was conducted (Guo et al., 2023). These studies represent significant milestones by providing reference genome assemblies for marine turtle studies. However, comprehensive population genomic studies are yet to be undertaken. Especially for hawksbill turtles, evaluating the genetic diversity of Indo-Pacific hawksbill turtle populations is imperative, given their critically endangered status and ecological position.

In this study, we analysed whole genome sequencing datasets obtained from a hawksbill turtle population nesting in Singapore, chosen as a representative population of Indo-Pacific marine turtles confronting substantial pressures from ongoing climate change and anthropogenic activities. Although most nesting beaches in Singapore are reclaimed land with narrow shoreline and considerable human activity, hawksbill turtles consistently nest there annually (Loh, 2019). The primary nesting sites in Singapore fall within regions expected to be significantly impacted by sea-level rise (Nguyen et al., 2022). The success rate of hatchlings at these Singapore nesting sites fluctuates due to adverse environmental factors, including light pollution from beachside lamp posts (Loh, 2019) and predation by monitor lizards and ghost crabs (Wong, 2019). Conservation strategies for this population nesting in Singapore are critical.

Our project’s objective is to assess the genetic diversity and demographic history of hawksbill turtles in Singapore using the whole genome sequencing datasets generated in this study. We have created a *de novo* genome assembly of the hawksbill turtle as the reference genome and performed high-quality whole genome sequencing on 35 samples, enabling a comprehensive population genetic study. This pioneering effort represents the first fine-resolution population genomic study for marine turtles, providing crucial insights for future research and conservation initiatives focused on hawksbill turtles.

## Results

### Maternal genetic diversity of hawksbill turtles in Singapore

Between 2018 to 2022, we collected unhatched eggs and unsuccessful hatchlings from 32 hawksbill turtle nests, and tissue samples from six adult hawksbill turtles in Singapore (Figure 1 and Supplementary table 1). Most of these nests (18 out of 32) were collected from the beach of East Coast Park that is narrow and reclaimed nature (Figure 1). Aside from Changi and East Coast Park beach, other nesting sites in the Southern Islands of Singapore, including Sentosa’s reclaimed beaches (3), Raffles Lighthouse (2), and Sisters’ Islands (3), which is a marine protected area. Four nests were relocated to the turtle hatchery on Sister’s Island due to unfavourable conditions at their original sites, such as human activity or risk of flooding. Samples from locations marked as unknown were collected during the COVID-19 outbreak’s circuit breaker lockdown in 2020, proper recording of their locations was not feasible. Additionally, samples from individual turtles were obtained from both live specimens and deceased ones found along the coast. In many cases, these turtles displayed injuries resulting from entanglement or were discovered washed ashore post mortem.

**Figure 1.**
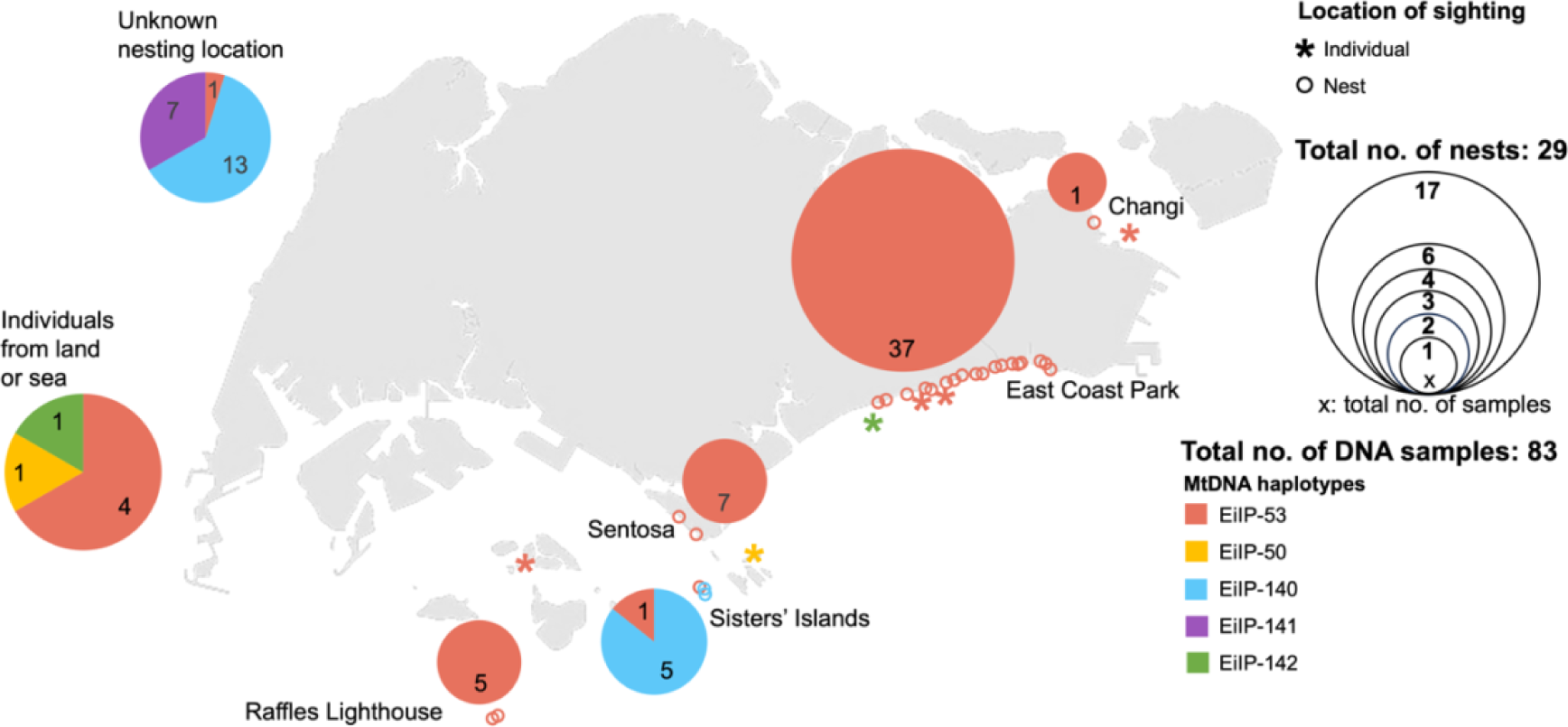
Distribution of hawksbill turtle nests we have collected for DNA samples and their mitochondrial haplotypes. The points marked on the map by (*) or (o) represent the locations where the hawksbill individual (live or dead) and nest are found respectively. Numbers within the pie charts (bottom) represent the number of DNA sample from each location, and the size of the pie charts represents the number of nests collected from each location (bolded in the legend). The colours of the pie charts represent the distribution of mitochondrial haplotypes in each location.

The availability of nesting samples collected depends on the accessibility of the nesting locations. Nests located in less accessible areas, such as Sisters’ Islands, often fully predated before they can be safeguarded for incubation and hatching. Consequently, Figure 1 does not offer a comprehensive representation the nesting distribution of hawksbill turtles in Singapore.

A total number of 83 DNA samples were sequenced, collected from 29 nests and 6 individual turtles. The number of DNA samples obtained from each nest varied between one and nine, depending on the hatchling success rate and DNA quality. The condition of the nests during sample collection significantly influenced the quality and yield of DNA. Out of the 83 extracted DNA samples, 69 samples, collected from 29 nests and four foraging and two individuals, were sequencing the mitochondrial (mt) control region, a widely used genetic marker in hawksbill turtle genetic studies (Abreu-Grobois et al., 2006). Among the 69 sequences, we identified five mt haplotypes: EiLP- 50, 53, EilP-140, 141, and 142. These haplotypes are designated by the Marine Mammal and Turtle Division of the National Oceanic and Atmospheric Administration (NOAA) based on their database of their hawksbill mt sequences. The most prevalent haplotypes among the five is EiIP-53, contributing exclusively to nests found in Changi, Tanah Merah, Sentosa, and Raffles Lighthouse in Singapore (Figure 1). Distinct haplotypes (EiIP-140 and EiIP-141) were found in Sisters’ Islands and the unknown location. Additionally, two other haplotypes (EiIP-50 and EiIP-142) were discovered in hawksbill individuals stranded either on land or at sea. The discovery of three novel haplotypes, EilP-140, 141, and 142, from two nesting and one foraging samples, showcases the significant genetic contribution of Singapore as nesting and foraging ground for the species.

The phylogenetic tree (Figure 2A) was constructed for the five distinctive haplotypes found in Singapore, combining them with 67 previously published mt haplotype sequences from the broader Indo-Pacific region (Bell & Jensen, 2018; Nishizawa et al., 2016; Vargas et al., 2016). All five Singapore haplotypes are clustered with Clade

**Figure 2.**
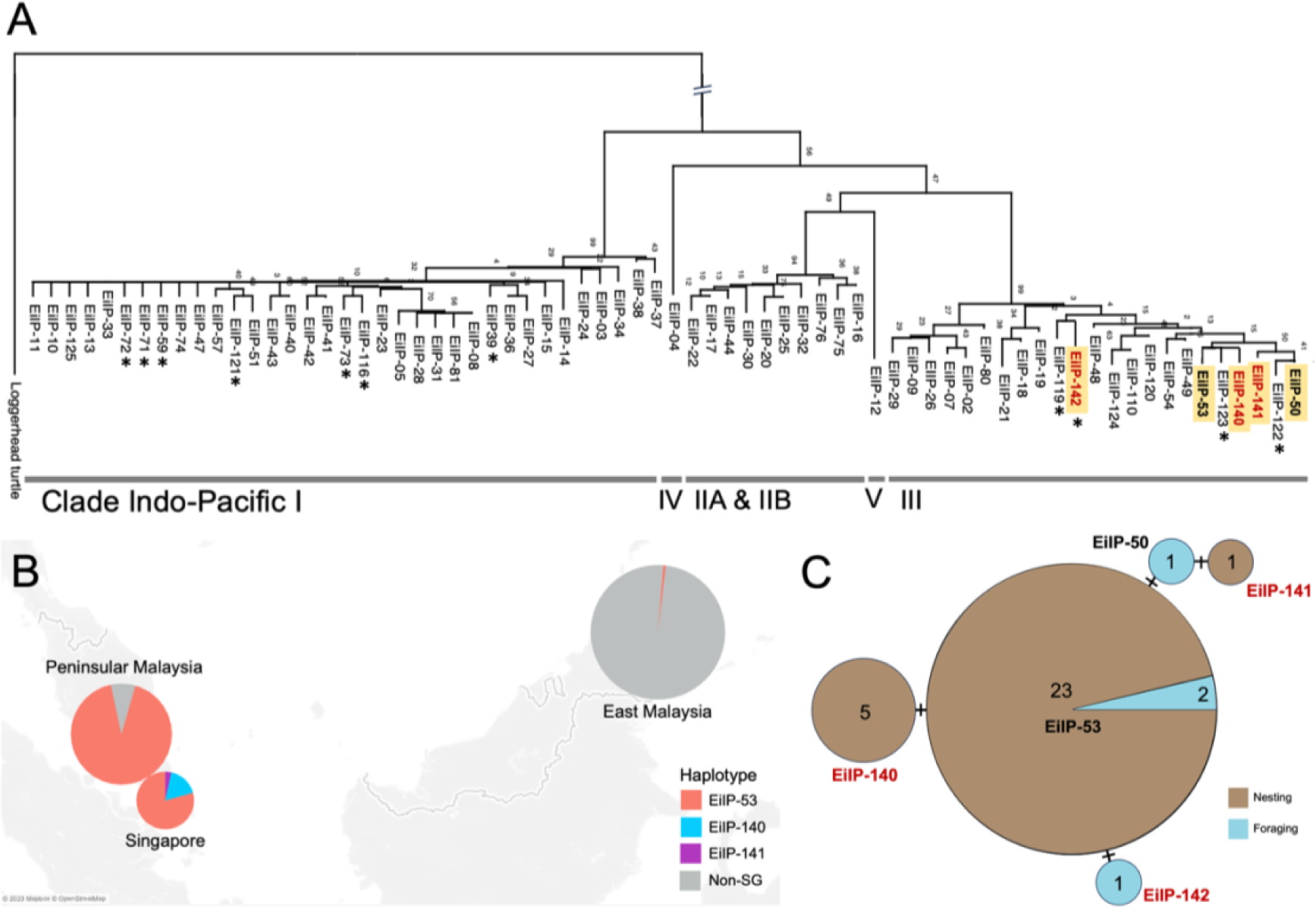
(A) Neighbour-joining phylogenetic tree of the haplotypes from Singapore (bolded and highlighted in yellow) and within the Indo-Pacific region, where all the Singapore haplotypes are clustered in Clade Indo-Pacific III. New haplotypes found in Singapore are written in red. Orphan haplotypes (only found in foraging sites) are marked with asterisks (*). **(B)** Nesting haplotype distribution and frequency within Southeast Asia (including Malaysia and Singapore), illustrating the overlap of Singapore haplotypes within this region. Haplotypes from East Malaysia is specifically found in Sabah Turtle Islands. **(C)** Network analysis of the genetic distribution and frequency of haplotypes per nest or individual found in Singapore.

Indo-Pacific III. Notably, three of these haplotypes (EiIP-140, EiIP-141, EiIP-142) discovered in Singapore are exclusive to this population as highlighted in red letters in Figure 2A. The nesting haplotype EiIP-53 is also present in Malay Peninsula (Pulau Redang, Melaka, Geliga and Johor) and East Malaysia (Sabah Turtle Islands) (Figure 2B) (refer to Table 3 in Supplementary for frequencies). These five Singapore haplotype sequences exhibit differences of only up to three nucleotide bases among them (Figure 2C). Previous studies (Arantes et al., 2020) have established genetic distinctions between the Atlantic and Eastern Pacific haplotypes, clustering separately from those in the Indo-Pacific. As a result, these Atlantic and Eastern Pacific haplotypes were not included in the phylogenetic tree.

### Whole genome assembly of hawksbill turtles

The *de novo* assembly of the hawksbill turtle genome was conducted to establish a reference dataset for constructing population whole genome sequencing datasets. The DNA sample used for this reference genome was extracted from the whole blood of a mother turtle, obtained after her attempt to lay eggs on the beach at East Coast Park. Employing the Oxford Nanopore PromethION and Illumina HiSeqX platforms, we assembled the hawksbill genome, designated as ECP20C, which comprises 2,490 contigs. The PromethION sequencing exhibited a depth of coverage at 57.9x, with an N50 of 3.4Mb (Supplementary Table 1). The estimated total genome size stands at 2.16 Gb, consistent with other marine turtle genome assemblies previously reported (Bentley et al., 2022; Guo et al., 2023). Conducting synteny analysis to compare the ECP20C genome assembly with other published hawksbill, green, and leatherback turtle genomes revealed a high degree of synteny across these species or assemblies. This indicates the completeness of our *de novo* genome assembly, which is sufficiently for a comparative genome analysis (Supplementary Figure 2).

### Whole genome diversity of hawksbill turtles in Singapore

We conducted high-depth sequencing on 35 samples using the Illumina HiseqX platform to explore the extent of genetic diversity within the population. The selection of samples for whole genome sequencing was determined based on the DNA quality and quantity. From these 35 whole genome datasets (Table 1), our analysis identified approximately 12 million single nucleotide polymorphisms (SNPs), with each genome containing between 4.5 to 5 million SNPs (Supplementary Fig 3). This level of resolution in genetic marker within whole genomes, in terms of number of SNPs, is significantly extensive, approximately 2.4 million times larger than that of the commonly used mt control region. Notably, even when compared to the entire mt genome, only 12 variants were identified among the 35 samples (Table 1), and six haplotypes were determined with the 12 variants.

**Table 1.** Comparison of genetic datasets of hawksbill turtle samples from Singapore.

|  | Mitochondria<br>I control<br>region | Mitochondrial<br>genome | Whole<br>genome |
| --- | --- | --- | --- |
| Length | 766 bp | 16,386 bp | 2.16 Gb |
| No. of samples | 69 | 35 | 35 |
| No. of nests and individuals | 30 nests<br>6 individuals | 20 nests<br>5 individuals | 20 nests<br>5 individuals |
| No. of biallelic single<br>nucleotide polymorphisms<br>(SNPs) | 5 | 12 | 11,960,448 |

We calculated identical-by-state (IBS) among all pairs of the 35 individual genomes to infer their genetic relatedness, using the whole genome sequencing datasets (Figure 3A). Out of the 35 samples, our findings revealed that 19 have at least a 3^rd^ degree relationship (first cousins) with one another, excluding those with 1^st^ degree relationships (parent-child and full sibling). This high level of relatedness observed among samples collected from various nests highlights the inbreeding and interconnected nature the hawksbill turtle population in Singapore.

**Figure 3.**
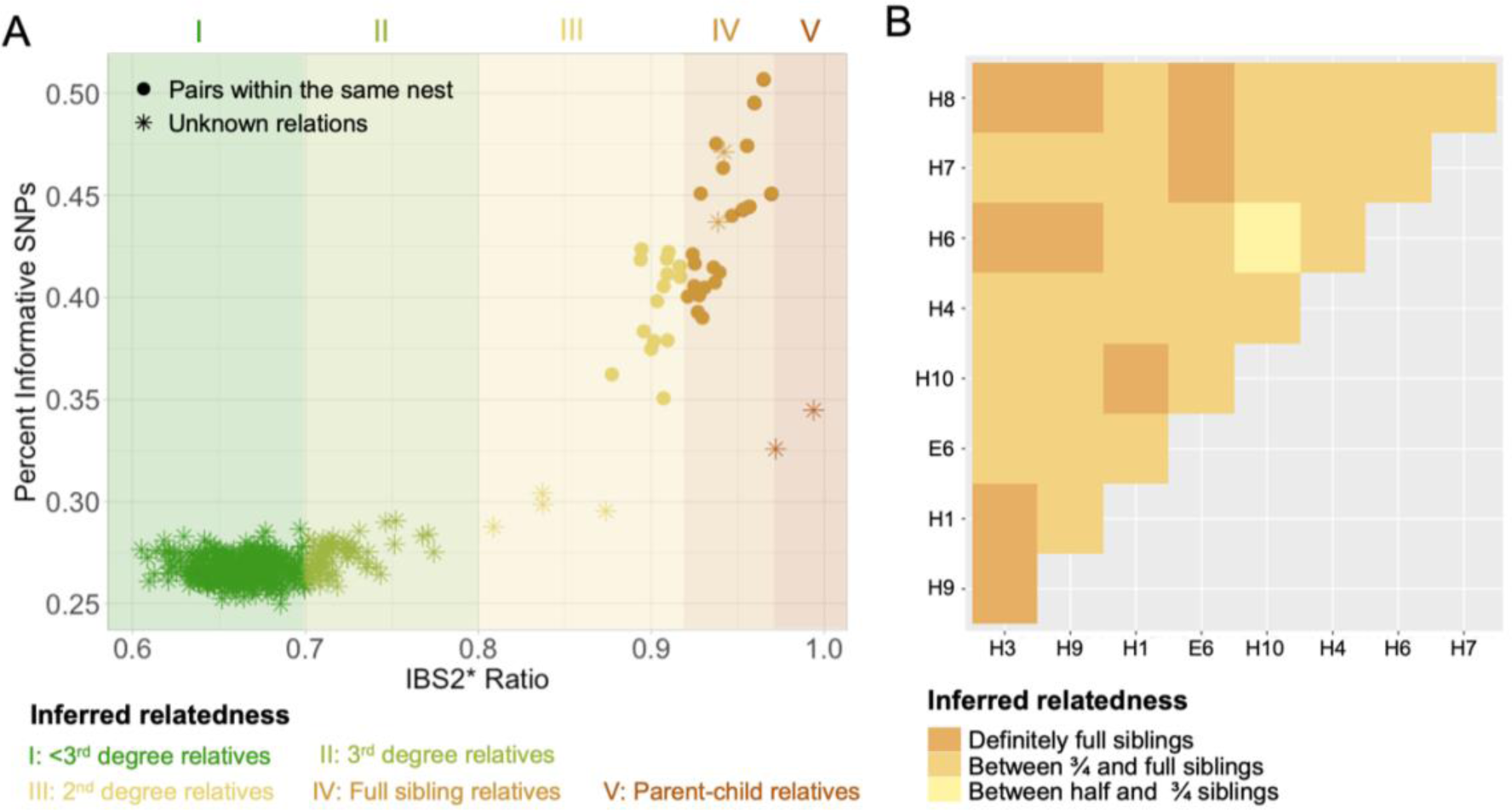
(A) Analysis from identity by state (IBS) suggests the presence of inbreeding as more than a third of the pairs exhibit at least a third-degree relationship (i.e. first cousins) with one another. The analysis also suggests the occurrence of multiple paternity involving closely related fathers within a single nest. **(B)** Identity by descent (IBD) plot of the siblings reveal the presence of ¾ siblings, suggesting that the fathers are closely related.

Moreover, our analysis identified two pairs of parent-child relationships, both of which were previously unknown. These pairs included a nest sample and a nesting mother individual which were collected in the same year.

Among the 35 samples, those collected from the same nest are considered as full siblings (solid dots in Figure 3A). The IBS analysis identified 23 pairs as full sibling relationships, including two pairs from different nests previously unidentified. Additionally, our analysis revealed four pairs of 2^nd^ degree relationships, such as aunts/uncles with nieces/nephews or half siblings.

An intriguing finding emerged from the nine samples collected from the same nest: some siblings show 2^nd^ degree relationships with one another. This suggests the possibility of multiple paternities contributing to this nest. However, despite this observation, clear identification of sibling clusters indicating multiple fathers among the nine samples is challenging based on both IBS and IBD analyses (Supplementary Figures 5 and 6). Many relationships among the nine genomes fall within the range between half and full siblings. The IBD within the nine siblings indicates that one pair is between three-quarter and half siblings, while the rest of the pairs are either full siblings or fall between full and three-quarter siblings (Figure 3B). This complexity in sibling relationships suggests the potential relatedness of the multiple parents. For example, if the fathers are siblings, their children with the same mother would be categorised as three-quarter siblings, sharing about 37.5% of their genome with each other. The IBD within the nine genomes supports the presence of multiple fathers and a mother who are closely related (Figure 3B).

### Demographic history of hawksbill turtles

To infer the population history of the hawksbill population, we conducted the multiple sequentially Markovian coalescence (MSMC) analysis (Figure 4). The MSMC estimates reveal a significant decline in the effective population size, from 40,000 to 5,000 which was an eight-fold reduction, occurring from 70,000 to 8,000 years ago. Simultaneously, there was a gradual increase in sea surface temperature (SST) and sea level rise (SLR) from 60,000 and 25,000 years ago, respectively, coinciding with the Last Glacial Maximum (LGM) and subsequent deglaciation period. Both SST and SLR started stabilising around 6,000 years ago, cooccurring with a slight increase in the effective population size of this hawksbill turtle population.

**Figure 4.**
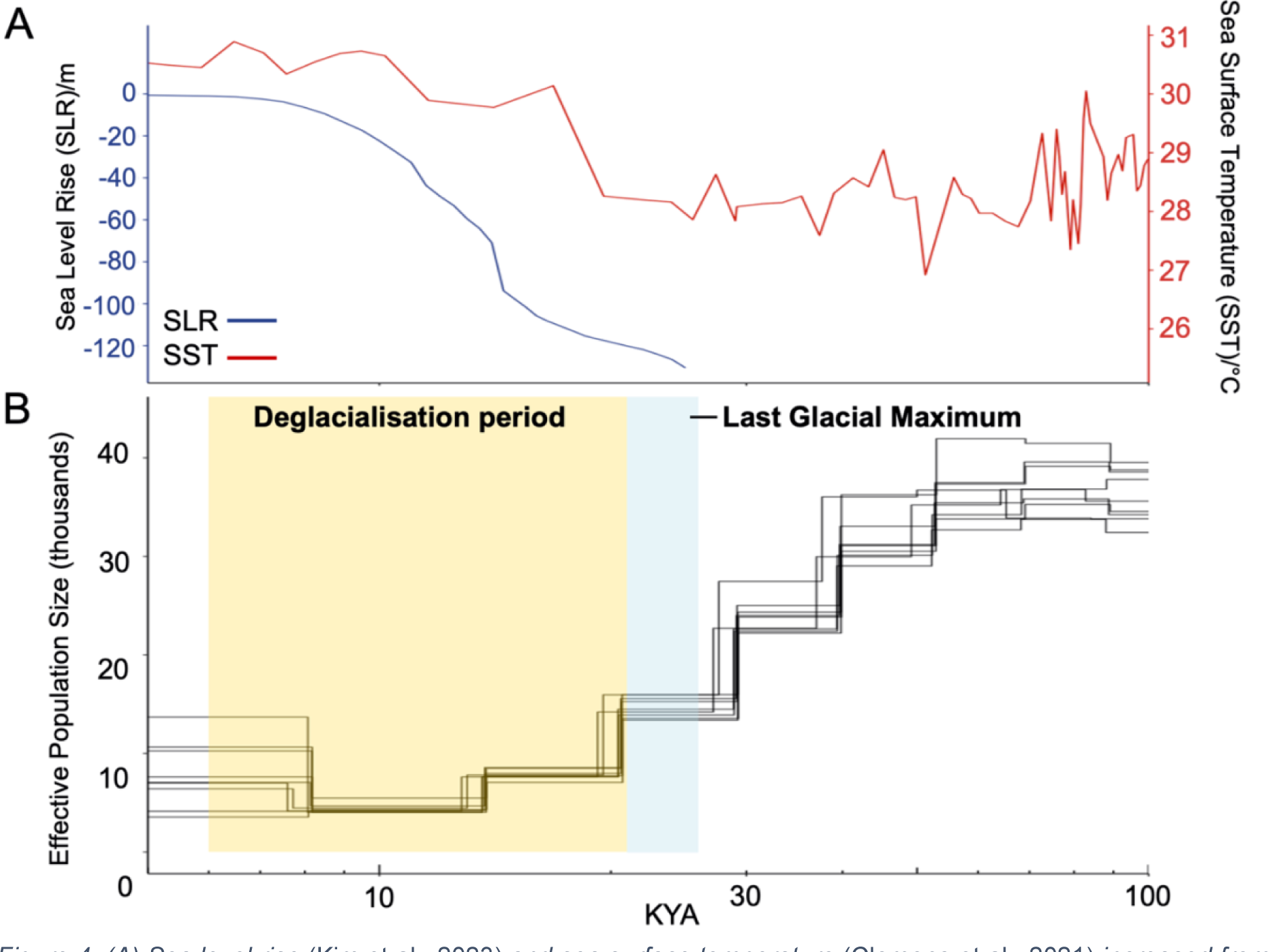
(A) Sea level rise (Kim et al., 2023) and sea surface temperature (Clemens et al., 2021) increased from 30 and 100 thousand years respectively. (B) Most recent effective population size decline occurred slightly prior and during the Last Glacial Maximum (LGM), calculated using Multiple sequentially Markovian coalescent (MSMC). A generation time of 35 years (Mortimer & Donnelly, 2008) and mutation rate of 2.5x10^-8^ mutations per nucleotide site per generation (Nachman & Crowell, 2000).

## Discussion

This study pioneered the generation of comprehensive whole genome population sequencing datasets and a *de novo* genome assembly of the hawksbill turtle, marking a significant milestone in this field. Using these datasets, we uncovered detailed insights into the genetic diversity and demographic history of a hawksbill turtle population nesting and foraging in Singapore.

Our findings underscore the pivotal role of whole genome sequencing datasets in enhancing the resolution of genetic markers for conservation genetic studies. The limitations inherent in biased and low-resolution genetic markers restrict the comprehensive examination of population genetic diversity and structure. For instances, our analysis based on mt haplotypes identified only five and six distinct maternal lineages based on the control region and whole mt genome, respectively, among hawksbill turtles in Singapore. With the most frequent haplotype, EilP-53, shared across various nesting sites in Malay Peninsula and Eastern Malaysia (Figure 2C), this mt marker could not differentiate Singapore nesting individuals from other populations. Effective genetic markers are crucial for tracing sample origins, which is particularly useful in combatting the illegal trade targeting hawksbill turtles’ shells.

Moreover, the usage of a genetic marker of mt haplotypes, representing maternal lineages, introduces bias and limits understanding of population structure. Our principal component analysis (PCA) using whole genome sequencing data illustrated the discordance between the mt control region and the nuclear genome (Supplementary Fig 4), emphasising the necessity for accurate population structure inference. This precision is crucial, not only for tracing sample origins but also for defining management units in conservation strategies for migratory species like marine turtles.

Using whole genome sequencing data from Singapore’s hawksbill turtle population, we conducted an unbiased and comprehensive examination of genetic diversity, revealing a mean of nucleotide diversity per site of 0.00138 in our dataset. This value positions the hawksbill turtle population between the reported values for leatherback turtle (0.000286) and green turtle (0.00246) by Bentley et al. (2022). Despite of hawksbill turtle’s past population decline due to prolonged environmental pressures (Figure 4), this level of genetic diversity surpasses that observed in leatherback turtles and even non-African human populations (Auton et al., 2015). This suggests the maintenance of genetic diversity from the large ancestral populations, indicating the potential to recovery with population growth.

Our analysis revealed the high degree of inbreeding within the Singapore hawksbill turtle population, supported by evidence of related parents within a multiple-paternity nest (Figure 3B and Supplementary Figures 5 & 6). While multiple paternity typically contributes to genetic diversity within a nest, our study showed close relatedness between parents, indicating significant levels of inbreeding. The continuation of inbreeding within a population could exponentially increase the genetic burden, potentially driving the population towards extinction.

Climate change emerges as a significant challenge for hawksbill turtles. Our inference of demographic history sheds light on how the population has suffered past extreme environmental changes. The decline in effective population size since before the last glacial maximum (LGM) continued through the last deglaciation until stabilisation of sea levels in the region. Notably, a slight population increase followed this stabilisation (Figure 4). This parallel occurrence highlights the profound impact of environmental changes on the hawksbill turtle population, indicating the potential for population recovery under favourable environmental conditions.

While our study clearly demonstrates the advantages of whole genome sequencing in marine turtle conservation genetics, its widespread application has encountered challenges due to the high cost of genome sequencing and data analysis. Nonetheless, as sequencing costs continue to decrease, the accumulation of whole genome sequencing data could facilitate future studies for the development of cost-effective genome-wide markers. This approach holds promise for substantially reducing expenses in fine-resolution conservation genetic study of marine turtles, making our study as the initial step towards this goal.

## Methods

### Sampling sites and sample collection

Nesting sites are distributed along the eastern and southern coasts of Singapore, including the Southern Islands of Singapore. Under the BIOME research permit issued by National Parks Board Singapore, dead hatchlings and eggs were collected post-hatchling emergence from 30 nests within nesting seasons of 2018 to 2022. Liver or blood samples were also extracted and collected from 5 stranded hawksbill turtles by the veterinary team or the Animal and Plant Health Centre of the National Parks Board, depending on its state of life, within the sampling period. All samples were transported in dry-ice and immediately stored at -20°C.

### DNA extraction

Dead hatchlings and eggs were dissected using sterilised disposable blades and 50 mg of liver tissues of hatchlings and partially developed fetuses, and whole unfertilised eggs were stored in sterile tubes for DNA extraction. Either 50 mg of liver or blood samples from stranded hawksbill turtles were also transferred to sterile tubes for DNA extraction.

DNA was extracted using QIAGEN’s Gentra Puregene Tissue Kit (QIAGEN, USA) for all liver and egg samples, following the protocol for tissue samples. For the blood samples, QIAGEN’s Blood and Cell Culture DNA Kit was used, following the protocol for blood samples and the Genomic-tip procedure (QIAGEN). The extracted DNA purity was determined using Nanodrop (Thermo Fisher Scientific, USA) and was quantified using the Qubit dsDNA HS assay kit (Invitrogen, USA). The samples were run on agarose gels to assess if the DNA was sheared.

Mitochondrial DNA amplification and analysis

Using polymerase chain reaction (PCR), the mitochondrial DNA (mtDNA) control region segment of approximately 800 base pairs was amplified for all DNA samples. The primer sequences are LCM15382 (5’-GCTTAACCCTAAAGCATTGG-3’) and H950g (5’-GTCTCGGATTTAGGGGTTTG-3’) (Abreu-Grobois et al., 2006). We followed the protocol mentioned in Abreu-Grobois et al. (2006). Subsequently, the PCR products were purified using the PureLink PCR Purification Kit (Invitrogen, USA). The mitochondrial haplotype sequences were obtained from the purified mtDNA samples using Sanger Sequencing.

The mitochondrial (mt) DNA sequences were aligned using MEGA X (Kumar et al., 2018), with ClustalW’s multiple sequence alignment program (Thompson et al., 1994). Sequences were trimmed to 766 base pairs and the number of nucleotide differences and distances between them were calculated. Subsequently, we visualised the results on a DNA network analysis plot using the ‘*haploNet’* function from the ‘*pegas*’ package in RStudio (RStudio Team, 2020).

MtDNA sequences reported by Nishizawa et al. (2016) and Vargas et al. (2016) were from the samples collected in Malaysia and the Indo-Pacific region, respectively, were downloaded from the public database (GenBank) (Supplementary Table 2). Nishizawa et al. (2016) collected hawksbill turtle samples from both nesting and foraging sites, while Vargas et al. (2016) sampled exclusively from nesting sites. Additionally, four more mtDNA haplotype sequences (EiIP-39, EiIP-59, EiIP-72 and EiIP-73), found in a foraging site in the Great Barrier Reef, Queensland, Australia (Bell & Jensen 2018), were retrieved from GenBank on NCBI.

Incorporating haplotype sequences found from foraging sites in the Indo-Pacific region can assist in the identification process if these foraging hawksbill turtles were indeed nesting in Singapore. A mtDNA of the loggerhead turtle (*Caretta caretta*) was also obtained from GenBank (GenBank accession number: AB830479.1) to serve as an outgroup for the construction of the phylogenetic tree. The loggerhead turtle was chosen because it is closest to the hawksbill turtle among the marine turtle species (Naro-Maciel et al., 2008).

Unique mtDNA control region haplotype sequences from Singapore, as well as 68 distinct haplotype sequences from Australia, Malaysia and the Indo-Pacific, along with the loggerhead turtle mtDNA sequence, were aligned by CLUSTALW with pairwise deletions and trimmed to 766 base pairs, based on shortest sequence in the dataset. Subsequently, using all the haplotypes in our dataset, a Neighbour-joining phylogenetic tree (Saitou & Nei, 1987) was constructed based on number of nucleotide differences using MEGA X (Kumar et al., 2018). A bootstrapping method of 500 replications by the number of nucleotide differences including transitions and transversions was used for the phylogenetic analysis.

### Whole genome sequencing and *de novo* genome assembly

High-molecular weight (HMW) DNA for *de novo* assembly was extracted from whole blood of an adult turtle. The extracted HMW DNA sample was used for preparing Oxford Nanopore (ONT) and Illumina libraries for sequencing. The ONT library was prepared following the manufacturer’s instructions for the SQK-LSK109 Genomic DNA by ligation library prep method and sequenced for 72 hours on the PromethION R9.4.1 flowcell (ONT, UK). The raw fast5 data was basecalled using Bonito v0.4.0 (https://github.com/nanoporetech/bonito) with the default model. The Illumina library was constructed according to Illumina’s TruSeq Nano DNA Sample Preparation protocol and sequenced on the Illumina HiSeq X at a read length of 150bp paired-end (Illumina, USA). The long reads generated from ONT sequencing was used to create the *de novo* genome assembly with Shasta 0.11.1 (Shafin et al., 2020), a *de novo* long-read assembler. Three rounds of Minimap2 (v2.17; Li, 2018) and Racon (v1.4.3; Vaser et al., 2017) were performed for read correction and contig polishing. Medaka 1.0.3 (https://github.com/nanoporetech/medaka) using neural networks to align individual sequencing reads against the assemblies. Lastly, the Illumina short-reads were used to polish the assembly using Hybrid Polisher (HyPo 1.0.3) (Kundu et al., 2019) to create the final genome assembly.

For generating population data, Illumina libraries were prepared for a total of 35 whole genomes of hawksbill turtle samples, including the sample for the *de novo* genome assembly, and sequenced on the Illumina HiSeqX platform, as previously described.

### Variant calling

Raw sequence reads generated from HiSeqX sequencing were quality checked using FastQC 0.11.8 (Andrews, 2010) and adapter-trimmed using Atropos 1.1.22 (Didion et al., 2017). The sequence reads were aligned using Burrows-Wheeler Alignment (BWA) 0.7.17 tool (Li & Durbin, 2009) to the hawksbill genome assembly we built in this study, and the 35 genomes have minimally 15x coverage depth. Variant calling was performed as per GATK 4.1.9.0’s (Van der Auwera & O’Connor, 2020) joint analysis workflow. GATK Haplotype Caller was used to call variants in each of the 35 genomes in gVCF mode (-ERC GVCF) to produce a record of genotype likelihoods and annotations at every site of the genome. GATK GenotypeGVCFs (Van der Auwera & O’Connor, 2020) was used for joint genotyping of the resulting gVCFs to identify variants in the dataset.

The total number of variants called from the 35 genomes are 16,772,988. For variant quality control, we removed variants with depth (DP) less than 350 and more than 1840 using VCFtools 0.1.17 (Danecek et al., 2011). The lower bound depth cut-off of 350 was determined by the number of samples (35) multiplied by a depth of 10 per sample. The upper bound cut-off was decided based on the third standard deviation, which is calculated to be about 1840. Singletons were also removed. The SNP genotype call rate of 5% or more were also filtered out. Indels and multiallelic variants were also removed. After these filtering, the total number of biallelic SNPs is 11,960,448, and we used this set of SNPs for downstream analyses.

The sequence reads of the 35 samples were also aligned to the reference sequence of the hawksbill mitochondrial genome (GenBank accession number: NC_12398.1) using BWA 0.7.17 tool (Li & Durbin, 2009) and mapped reads were extracted. Variant calling was performed using BCFtools (Li, 2011) and variants with depth (DP) less than 20 were removed. The variants were then validated against their aligned reads using Integrative Genomics Viewer (IGV) (Robinson et al., 2011).

### Population genomic analysis

Nucleotide diversity was calculated to estimate the extent of genetic diversity of a population for 21 unrelated genomes. Nucleotide diversity was calculated using VCFtools 0.1.17 (Danecek et al., 2011) with the option “--window-pi” using a window of 10000 bp, and the mean was calculated.

PLINK 1.90 beta (Chang et al., 2015) was used to generate the eigen files for principal component analysis (PCA) using the “--pca” command. The PCA results of 21 genomes excluding samples with 1^st^ degree relationships were then visualised on R Studio 2022.07.1 Build 554 (RStudio Team, 2020) using the “ggplot2” package. 21 unrelated genomes after removing related and outlier samples based on high homozygosity. The genetic relationships between the individuals were inferred using identity by descent (IBD) and identity by state (IBS2) for 31 genomes after removing highly homozygous genomes. IBS counts were calculated using by SNPduo (Roberson & Pevsner, 2009) with the option “--counts”. The IBS2*_ratio was calculated based on the ratio of IBS2*/(IBS0+IBS2*). The percent informative SNPs are then calculated by the formula (IBS0+IBS2*)/(IBS0+IBS1+IBS2) (Stevens et al., 2011) and visualised on RStudio 2022.07.1 Build 554 (RStudio Team, 2020) using “ggplot2” package with the percent informative SNPs as the y-axis values and IBS2*_ratio as the x-axis values. IBD was estimated on PLINK 1.90 beta (Chang et al., 2015) by calculating PI_HAT values for each pair of individuals, based on the formula PI_HAT = P(IBD=2)+0.5*P(IBD=1), where IBD=1 and IBD=2 represent have 1 or 2 alleles over the loci. The results were visualised as a heatmap on RStudio (RStudio Team, 2020) using “ggplot2” package.

Unique family configurations of 35 genomes were also estimated using KING 2.3.1 (Manichaikul et al., 2010) with the “--related” option on IBD segments. VCFtools (Danecek et al., 2011) and PLINK 1.90 beta (Chang et al., 2015) were used to generate the input data in the required format for KING. The options “--degree 3” and “--rplot” was used alongside in KING to generate all family configurations up to 2^nd^ degree relationships.

Each contig of the 35 genomes were phased using SHAPEIT4 (Delaneau et al., 2019) with no reference. 19 contigs with no variants were removed and a total 2471 contigs were phased. The phased genome datasets were used for the MSMC2 demographic inference.

Demographic inference was carried out using multiple sequentially Markovian coalescent (MSMC2) (Schiffels & Durbin, 2020) to infer effective population size change over time. 18 unrelated and high-quality genomes were selected for the MSMC analysis. The analysis was performed for every pair of the samples. The estimates were scaled with a generation time of 35 years (Mortimer & Donnelly, 2008) and mutation rate of 2.5 x 10^-8^ /site/generation (Nachman & Crowell, 2000). The MSMC results were visualised on RStudio (RStudio Team, 2020). Environmental data including sea level rise and sea surface temperature data from (Kim et al., 2023) and (Clemens et al., 2021) respectively were visualised with the same temporal period.

