## Supplementary for "Population genomic analysis reveals high inbreeding in a Hawksbill turtle population nesting in Singapore"

*Table 1. Metadata of hawksbill turtle samples from Singapore used in this study.*

| DNA Sample | Location | Nest ID/ Foraging | Year | Generated sequencing data | Mt haplotype |
|---|---|---|---|---|---|
| ECP20B | At sea | Foraging | 2020 | Whole genome | EiIP-53 |
| ECP20C | At sea | Foraging | 2020 | Whole genome | EiIP-53 |
| RHB20 | At sea | Foraging | 2020 | Both | EiIP-50 |
| PH21D | At sea | Foraging | 2021 | Mitochondria | EiIP-53 |
| ECP21DJ | At sea | Foraging | 2021 | Both | EiIP-142 |
| CBP22A | At sea | Foraging | 2022 | Both | EiIP-53 |
| O18F1 | Unknown | O18 | 2018 | Both | EiIP-53 |
| ECP19AE1 | East Coast Park | ECP19A | 2019 | Mitochondria | EiIP-53 |
| ECP19AE9 | East Coast Park | ECP19A | 2019 | Both | EiIP-53 |
| ECP19BH1 | East Coast Park | ECP19B | 2019 | Both | EiIP-53 |
| ECP19CH1 | East Coast Park | ECP19C | 2019 | Both | EiIP-53 |
| SEN19AE5 | Sentosa | SEN19A | 2019 | Mitochondria | EiIP-53 |
| SEN19AE12 | Sentosa | SEN19A | 2019 | Mitochondria | EiIP-53 |
| SEN19AE22 | Sentosa | SEN19A | 2019 | Both | EiIP-53 |
| SEN19AE24 | Sentosa | SEN19A | 2019 | Mitochondria | EiIP-53 |
| SIS19AE1 | Sisters' Islands | SIS19A | 2019 | Mitochondria | EiIP-140 |
| SIS19BE1 | Sisters' Islands | SIS19B | 2019 | Mitochondria | EiIP-140 |
| SIS19BE2 | Sisters' Islands | SIS19B | 2019 | Mitochondria | EiIP-140 |
| SIS19BE3 | Sisters' Islands | SIS19B | 2019 | Mitochondria | EiIP-140 |
| SIS19BE4 | Sisters' Islands | SIS19B | 2019 | Mitochondria | EiIP-140 |
| LOC19AH1 | Unknown | LOC19A | 2019 | Mitochondria | EiIP-141 |
| LOC19AH2 | Unknown | LOC19A | 2019 | Mitochondria | EiIP-141 |
| LOC19AH3 | Unknown | LOC19A | 2019 | Mitochondria | EiIP-141 |
| LOC19AH4 | Unknown | LOC19A | 2019 | Mitochondria | EiIP-141 |
| LOC19AH5 | Unknown | LOC19A | 2019 | Mitochondria | EiIP-141 |
| LOC19AH6 | Unknown | LOC19A | 2019 | Mitochondria | EiIP-141 |
| LOC19AH7 | Unknown | LOC19A | 2019 | Both | EiIP-141 |
| LOC19CEH1 | Unknown | LOC19CE | 2019 | Whole genome | EiIP-140 |
| LOC19CEH2 | Unknown | LOC19CE | 2019 | Mitochondria | EiIP-140 |
| LOC19CEH3 | Unknown | LOC19CE | 2019 | Whole genome | EiIP-140 |
| LOC19CEH4 | Unknown | LOC19CE | 2019 | Whole genome | EiIP-140 |
| LOC19CEH6 | Unknown | LOC19CE | 2019 | Whole genome | EiIP-140 |
| LOC19CEH7 | Unknown | LOC19CE | 2019 | Whole genome | EiIP-140 |
| LOC19CEH8 | Unknown | LOC19CE | 2019 | Whole genome | EiIP-140 |
| LOC19CEH9 | Unknown | LOC19CE | 2019 | Whole genome | EiIP-140 |

| | | | | | |
|---|---|---|---|---|---|
| LOC19CEH10 | Unknown | LOC19CE | 2019 | Whole genome | EiIP-140 |
| LOC19CEE3 | Unknown | LOC19CE | 2019 | Mitochondria | EiIP-140 |
| LOC19CEE5 | Unknown | LOC19CE | 2019 | Mitochondria | EiIP-140 |
| LOC19CEE6 | Unknown | LOC19CE | 2019 | Both | EiIP-140 |
| LOC19DH1 | Unknown | LOC19D | 2019 | Mitochondria | EiIP-140 |
| ECP20AH1 | East Coast Park | ECP20A | 2020 | Both | EiIP-53 |
| ECP20AE1 | East Coast Park | ECP20A | 2020 | Mitochondria | EiIP-53 |
| ECP20AE2 | East Coast Park | ECP20A | 2020 | Mitochondria | EiIP-53 |
| ECP20AE3 | East Coast Park | ECP20A | 2020 | Mitochondria | EiIP-53 |
| ECP20AE8 | East Coast Park | ECP20A | 2020 | Mitochondria | EiIP-53 |
| ECP20BE1 | East Coast Park | ECP20B | 2020 | Both | EiIP-53 |
| ECP20BE2 | East Coast Park | ECP20B | 2020 | Mitochondria | EiIP-53 |
| ECP20DE1 | East Coast Park | ECP20D | 2019 | Mitochondria | EiIP-53 |
| ECP20DE2 | East Coast Park | ECP20D | 2020 | Whole genome | EiIP-53 |
| ECP20EH1 | East Coast Park | ECP20E | 2020 | Mitochondria | EiIP-53 |
| ECP20EH2 | East Coast Park | ECP20E | 2020 | Both | EiIP-53 |
| ECP20FH2 | East Coast Park | ECP20F | 2020 | Whole genome | EiIP-53 |
| ECP20FE2 | East Coast Park | ECP20F | 2021 | Mitochondria | EiIP-53 |
| ECP20GE1 | East Coast Park | ECP20G | 2020 | Both | EiIP-53 |
| ECP20GE2 | East Coast Park | ECP20G | 2020 | Mitochondria | EiIP-53 |
| ECP20HE1 | East Coast Park | ECP20H | 2020 | Mitochondria | EiIP-53 |
| ECP20HH1 | East Coast Park | ECP20H | 2020 | Mitochondria | EiIP-53 |
| ECP20HH2 | East Coast Park | ECP20H | 2020 | Both | EiIP-53 |
| ECP20IE1 | East Coast Park | ECP20I | 2020 | Mitochondria | EiIP-53 |
| ECP20IE2 | East Coast Park | ECP20I | 2020 | Both | EiIP-53 |
| ECP20JE1 | East Coast Park | ECP20J | 2020 | Both | EiIP-53 |
| ECP20JE2 | East Coast Park | ECP20J | 2020 | Mitochondria | EiIP-53 |
| RLH20AE1 | Raffles Lighthouse | RLH20A | 2020 | Mitochondria | EiIP-53 |
| RLH20AE2 | Raffles Lighthouse | RLH20A | 2020 | Mitochondria | EiIP-53 |
| RLH20BE1 | Raffles Lighthouse | RLH20B | 2020 | Mitochondria | EiIP-53 |
| RLH20BE2 | Raffles Lighthouse | RLH20B | 2020 | Mitochondria | EiIP-53 |
| RLH20BE4 | Raffles Lighthouse | RLH20B | 2020 | Whole genome | EiIP-53 |
| TM20AE9 | East Coast Park | TM20A | 2020 | Whole genome | EiIP-53 |
| TM20AE14 | East Coast Park | TM20A | 2020 | Mitochondria | EiIP-53 |
| CH21AH1 | Changi | CH21A | 2021 | Both | EiIP-53 |
| BS21AH1 | Sisters' Islands | BS21A | 2021 | Both | EiIP-53 |
| ECP21CE2 | East Coast Park | ECP21C | 2021 | Both | EiIP-53 |
| ECP21CE4 | East Coast Park | ECP21C | 2021 | Mitochondria | EiIP-53 |
| ECP21CE5 | East Coast Park | ECP21C | 2021 | Mitochondria | EiIP-53 |
| SEN21BE2 | Sentosa | SEN21B | 2021 | Mitochondria | EiIP-53 |
| SEN21BE3 | Sentosa | SEN21B | 2021 | Mitochondria | EiIP-53 |

| | | | | | |
|---|---|---|---|---|---|
| SEN21BE4 | Sentosa | SEN21B | 2021 | Mitochondria | EiIP-53 |
| ECP22BE1 | East Coast Park | ECP22B | 2022 | Mitochondria | EiIP-53 |
| ECP22CH1 | East Coast Park | ECP22C | 2022 | Both | EiIP-53 |
| TM22BH1 | East Coast Park | TM22B | 2022 | Both | EiIP-53 |
| TM22BH2 | East Coast Park | TM22B | 2022 | Both | EiIP-53 |
| TM22BH3 | East Coast Park | TM22B | 2022 | Both | EiIP-53 |
| TM22BH4 | East Coast Park | TM22B | 2022 | Both | EiIP-53 |

*Table 2. Mitochondrial haplotypes from the Indo-Pacific region used in Figure 2A.*

| Haplotype | GenBank # | Location | Nesting/ Foraging | Reference |
|---|---|---|---|---|
| EiIP-02 | KT934050 | Milman Island, north Queensland, Australia | Nesting | Vargas et al. (2016) |
| EiIP-03 | KT934051 | Solomon Islands | Nesting | Vargas et al. (2016) |
| EiIP-04 | KT934052 | Milman Island, north Queensland, Australia | Nesting | Vargas et al. (2016) |
| EiIP-05 | KT934053 | Milman Island, north Queensland, Australia | Nesting | Vargas et al. (2016) |
| EiIP-07 | KT934054 | Milman Island, north Queensland, Australia | Nesting | Vargas et al. (2016) |
| EiIP-08 | KT934055 | Milman Island, north Queensland; Western Australia; northeast Arnhem Land, Northern Territory, Australia (N); Melaka (F) | Nesting, Foraging | Vargas et al. (2016), Nishizawa et al., (2016) |
| EiIP-09 | KT934056 | Milman Island, north Queensland; northeast Arnhem Land, Northern Territory, Australia | Nesting | Vargas et al. (2016) |
| EiIP-10 | KT934057 | Iran Northwest | Nesting | Vargas et al. (2016) |
| EiIP-11 | KT934058 | Iran Northwest | Nesting | Vargas et al. (2016) |
| EiIP-12 | KT934059 | Iran Southeast | Nesting | Vargas et al. (2016) |
| EiIP-13 | KT934060 | Iran Southeast | Nesting | Vargas et al. (2016) |
| EiIP-14 | KT934061 | Iran Southeast | Nesting | Vargas et al. (2016) |
| EiIP-15 | KT934062 | Iran Southeast | Nesting | Vargas et al. (2016) |
| EiIP-16 | KT934063 | Amirantes Islands; Platte Island; Granitics Islands; Chagos Archipelago, Seychelles | Nesting | Vargas et al. (2016) |
| EiIP-17 | KT934064 | Amirantes Islands; Platte Island; Granitics Islands; Chagos Archipelago, Seychelles | Nesting | Vargas et al. (2016) |
| EiIP-18 | KT934065 | Aldabra Group; Amirantes Islands; Platte Island; Granitics Islands; Chagos Archipelago, Seychelles | Nesting | Vargas et al. (2016) |
| EiIP-19 | KT934066 | Amirantes Islands; Platte Island; Granitics Islands, Seychelles | Nesting | Vargas et al. (2016) |
| EiIP-20 | KT934067 | Platte Island, Seychelles | Nesting | Vargas et al. (2016) |
| EiIP-21 | KT934068 | Aldabra Group; Amirantes Islands, Seychelles | Nesting | Vargas et al. (2016) |
| EiIP-22 | KT934069 | Granitics Islands, Seychelles | Nesting | Vargas et al. (2016) |
| EiIP-23 | KT934070 | Solomon Islands | Nesting | Vargas et al. (2016) |
| EiIP-24 | KT934071 | Solomon Islands | Nesting | Vargas et al. (2016) |

| | | | | |
|---|---|---|---|---|
| EiIP-25 | KT934072 | Granitics Islands, Seychelles | Nesting | Vargas et al. (2016) |
| EiIP-26 | KT934073 | northeast Arnhem Land, Northern Territory, Australia | Nesting | Vargas et al. (2016) |
| EiIP-27 | KT934074 | Saudi Arabia | Nesting | Vargas et al. (2016) |
| EiIP-28 | KT934075 | Western Australia | Nesting | Vargas et al. (2016) |
| EiIP-29 | KT934076 | Milman Island, north Queensland, Australia | Nesting | Vargas et al. (2016) |
| EiIP-30 | KT934077 | Granitics Islands, Seychelles | Nesting | Vargas et al. (2016) |
| EiIP-31 | KT934078 | Milman Island, north Queensland; northeast Arnhem Land, Northern Territory, Australia | Nesting | Vargas et al. (2016) |
| EiIP-32 | KT934079 | Chagos Archipelago, Seychelles | Nesting | Vargas et al. (2016) |
| EiIP-33 | KT934080 | Iran Northwest; Iran Southwest; Saudi Arabia; Aldabra Group; Platte Island; Granitics Islands, Seychelles; Milman Island, north Queensland, Australia; Solomon Islands (N)<br>Tun Sakaran Marine Park; Pulau Sipadan; Melaka, Malaysia (F) | Nesting | Vargas et al. (2016), Nishizawa et al., (2016) |
| EiIP-34 | KT934081 | Solomon Islands | Nesting | Vargas et al. (2016) |
| EiIP-36 | KT934082 | Iran Northwest; Iran Southwest; Saudi Arabia | Nesting | Vargas et al. (2016) |
| EiIP-37 | KT934083 | Western Australia | Nesting | Vargas et al. (2016) |
| EiIP-38 | KT934084 | Western Australia | Nesting | Vargas et al. (2016) |
| EiIP-40 | KT934085 | Iran Southwest | Nesting | Vargas et al. (2016) |
| EiIP-41 | KT934086 | Iran Northwest; Iran Southwest | Nesting | Vargas et al. (2016) |
| EiIP-42 | KT934087 | Iran Northwest | Nesting | Vargas et al. (2016) |
| EiIP-43 | KT934088 | Solomon Islands | Nesting | Vargas et al. (2016) |
| EiIP-44 | KT934089 | Chagos Archipelago, Seychelles | Nesting | Vargas et al. (2016) |
| EiIP-47 | KT934090 | East Malaysia (N); Pulau Sipadan (F) | Nesting, Foraging | Vargas et al. (2016), Nishizawa et al., (2016) |
| EiIP-48 | KT934091 | East Malaysia | Nesting | Vargas et al. (2016), Nishizawa et al., (2016) |
| EiIP-49 | KT934092 | East Malaysia; Peninsular Malaysia | Nesting | Vargas et al. (2016), Nishizawa et al., (2016) |
| EiIP-50 | KT934093 | East Malaysia | Nesting | Vargas et al. (2016) |
| EiIP-51 | KT934094 | East Malaysia | Nesting | Vargas et al. (2016), Nishizawa et al., (2016) |
| EiIP-53 | KT934095 | East Malaysia; Peninsular Malaysia | Nesting | Vargas et al. (2016), Nishizawa et al., (2016) |
| EiIP-54 | KT934096 | Peninsular Malaysia | Nesting | Vargas et al. (2016), Nishizawa et al., (2016) |
| EiIP-57 | KT934097 | Milman Island, north Queensland, Australia | Nesting | Vargas et al. (2016) |

| | | | | |
|---|---|---|---|---|
| EiIP-75 | KT934098 | Granitics Islands, Seychelles | Nesting | Vargas et al. (2016) |
| EiIP-76 | KT934099 | Amirantes Islands, Seychelles | Nesting | Vargas et al. (2016) |
| EiIP-80 | KT934100 | northeast Arnhem Land, Northern Territory, Australia | Nesting | Vargas et al. (2016) |
| EiIP-81 | KT934101 | northeast Arnhem Land, Northern Territory, Australia | Nesting | Vargas et al. (2016) |
| EiIP-110 | KT072791 | Peninsular Malaysia | Nesting | Nishizawa et al., (2016) |
| EiIP-124 | KR706175 | Peninsular Malaysia | Nesting | Nishizawa et al., (2016) |
| EiIP-125 | KR706176 | East Malaysia (N); Pulau Sipadan (F) | Nesting, Foraging | Vargas et al. (2016), Nishizawa et al., (2016) |
| EiIP-39 | KT964291 | Great Barrier Reef, Queensland, Australia | Foraging | Bell & Jensen (2018) |
| EiIP-59 | KT964292 | Great Barrier Reef, Queensland, Australia | Foraging | Bell & Jensen (2018) |
| EiIP-71 | KT964293 | Pulau Sipadan | Foraging | Nishizawa et al., (2016) |
| EiIP-72 | KT964294 | Great Barrier Reef, Queensland, Australia | Foraging | Bell & Jensen (2018) |
| EiIP-73 | KT964295 | Great Barrier Reef, Queensland, Australia | Foraging | Bell & Jensen (2018) |
| EiIP-74 | KT964296 | Tun Sakaran Marine Park | Foraging | Nishizawa et al., (2016) |
| EiIP-116 | KT072796 | Tun Sakaran Marine Park | Foraging | Nishizawa et al., (2016) |
| EiIP-119 | KR706170 | Tun Sakaran Marine Park | Foraging | Nishizawa et al., (2016) |
| EiIP-120 | KR706171 | Tun Sakaran Marine Park | Foraging | Nishizawa et al., (2016) |
| EiIP-121 | KR706172 | Pulau Sipadan | Foraging | Nishizawa et al., (2016) |
| EiIP-122 | KR706173 | Pulau Tiga | Foraging | Nishizawa et al., (2016) |
| EiIP-123 | KR706174 | Pulau Tiga | Foraging | Nishizawa et al., (2016) |

*Table 3. Mitochondrial haplotype frequencies from Southeast Asia used in Figure 2B.*

| Region | Nesting locations | **EiIP49** | **EiIP53** | **EiIP54** | **EiIP110** | **EiIP124** | **EiIP17** | **EiIP47** | **EiIP48** | **EiIP50** | **EiIP51** | **EiIP125** | **EiIP140** | **EIIP141** |
|---|---|---|---|---|---|---|---|---|---|---|---|---|---|---|
| Peninsular Malaysia | Pulau Redang, Melaka, Geliga, Johor | 1 | 82 | 2 | 2 | 2 | 0 | 0 | 0 | 0 | 0 | 0 | 0 | 0 |
| East Malaysia | Sabah Turtle Islands | 43 | 1 | 0 | 0 | 0 | 1 | 61 | 45 | 1 | 4 | 1 | 0 | 0 |
| Singapore | Singapore | 0 | 23 | 0 | 0 | 0 | 0 | 0 | 0 | 0 | 0 | 0 | 5 | 1 |

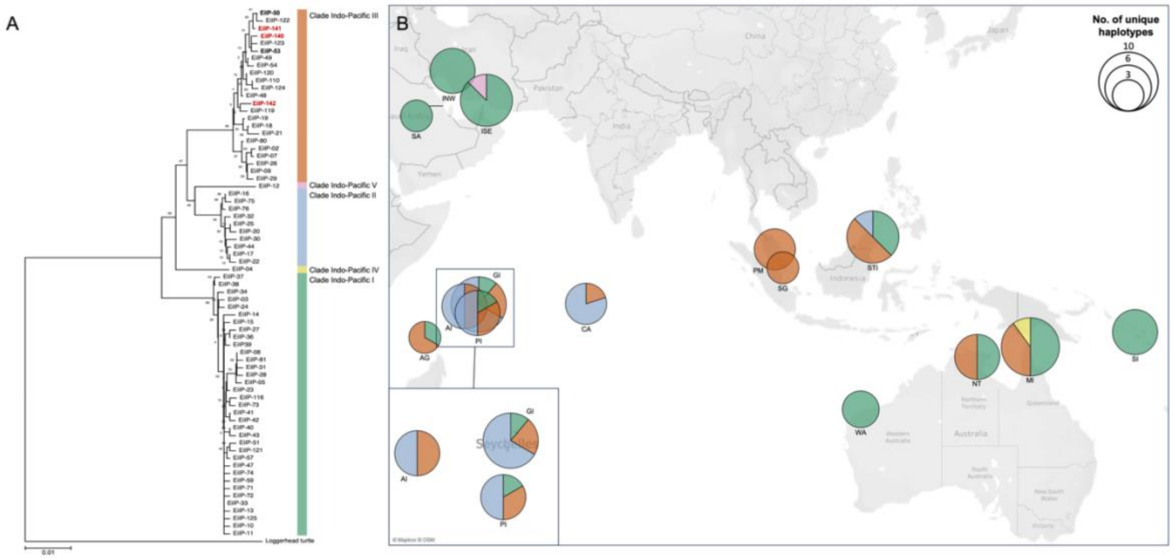


*Figure 1. (A) Neighbour-joining tree of the haplotypes from Singapore (bolded) and within the Indo-Pacific region, where all the Singapore haplotypes are clustered in Clade Indo-Pacific III. New haplotypes found in Singapore are written in red. (B) Nesting haplotype distribution and frequency within the Indo-Pacific region, illustrating the overlap of Singapore haplotypes within this region. Abbreviations are: Iran Northwest (INW); Iran Southeast (ISE); Saudi Arabia (SA); Aldabra Group, Seychelles (AG); Amirantes Islands, Seychelles (AI); Platte Island, Seychelles (PI); Granitics Islands (GI); Chagos Archipelago (CA); Sabah Turtle Islands, East Malaysia (STI); Peninsular Malaysia (PM); Singapore (SG); Rosemary and Varanus Islands, Western Australia (WA); northeast Arnhem Land, Northern Territory (NT); Milman Island, north Queensland (MI); Solomon Islands (SI).*

*Table 4. Hawksbill genome assembly statistics.*

| | |
|---|---|
| No. of contigs | 2490 |
| Largest contig (Mb) | 24.2 |
| N50 (Mb) | 3.4 |
| GC (%) | 44.0 |
| Coverage Depth (fold) | 57.9 |
| Genome length (Gb) | 2.16 |

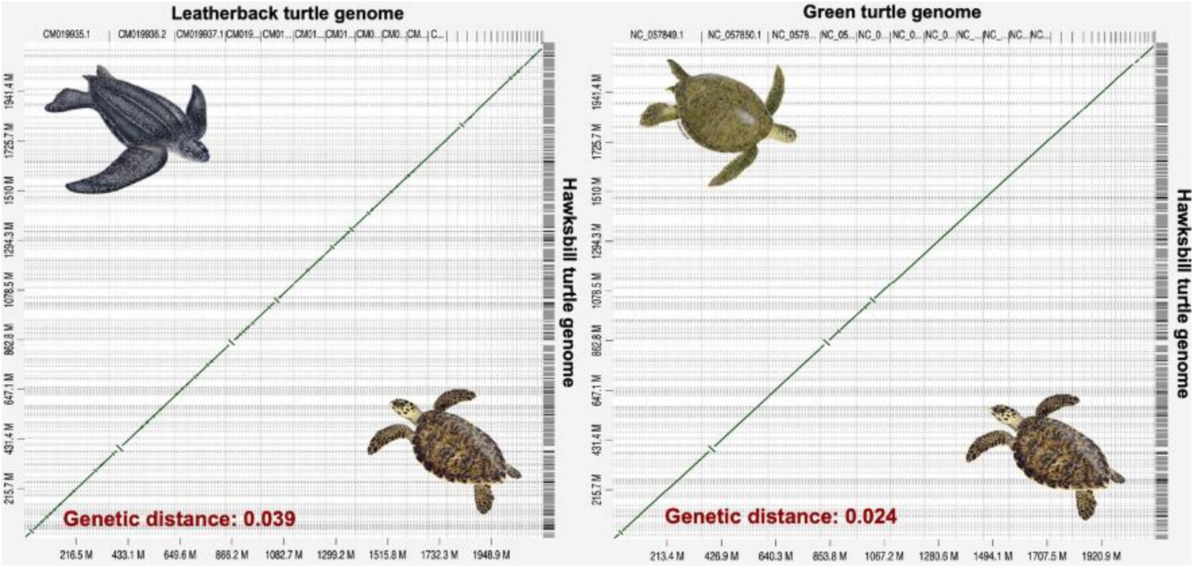


*Figure 2. Genome comparisons between our hawksbill genome assembly with the published genome assemblies of the green and leatherback turtles using synteny. The genetic distances were also calculated using number of SNPs divided by the total length.*

The D-GENIES synteny plots provide clear evidence that the hawksbill turtle genome exhibits a high degree of synteny with the green turtle genome, with only a few minor inversions (Figure 2). These regions of inversions were identified using the annotations of the green genome assembly, aiming to gain a better understanding of interspecies differences with respect to the hawksbill turtle.

A total of seven genes were disrupted at these boundaries and only two of them had no known homologs within the hawksbill turtle genome. One of these two genes is CUB and Sushi multiple domains 3 gene (CSMD3). CSMD3, found in chromosome 4 of the green turtle genome, is a regulator of dendrite development and mutations of this gene have been frequently identified in patients with schizophrenia and autism (Mizukami et al., 2016). The second disrupted gene, found in chromosome 5 of the green turtle genome, is phosphodiesterase 4D and it controls the availability of cyclic adenosine monophosphate (cAMP), particularly for memory formation (Ricciarelli & Fedele, 2015).

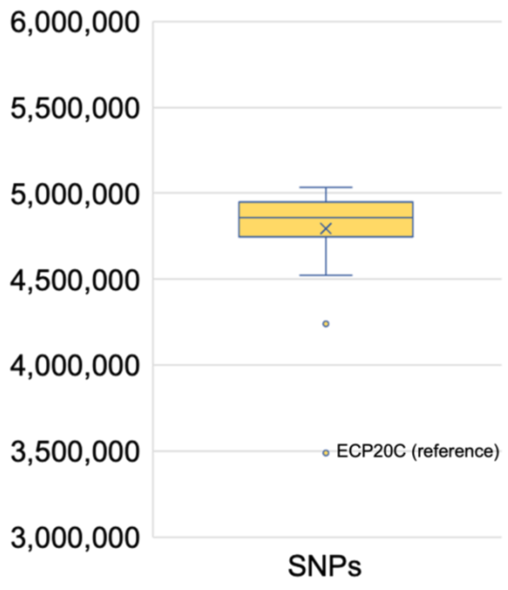


*Figure 3. Number of single nucleotide polymorphisms (SNPs) between each sample and our hawksbill reference assembly. There are approximately 4.5-5 million SNPs between hawksbill individuals, which is more than human individuals globally (3-4 million SNPs).*

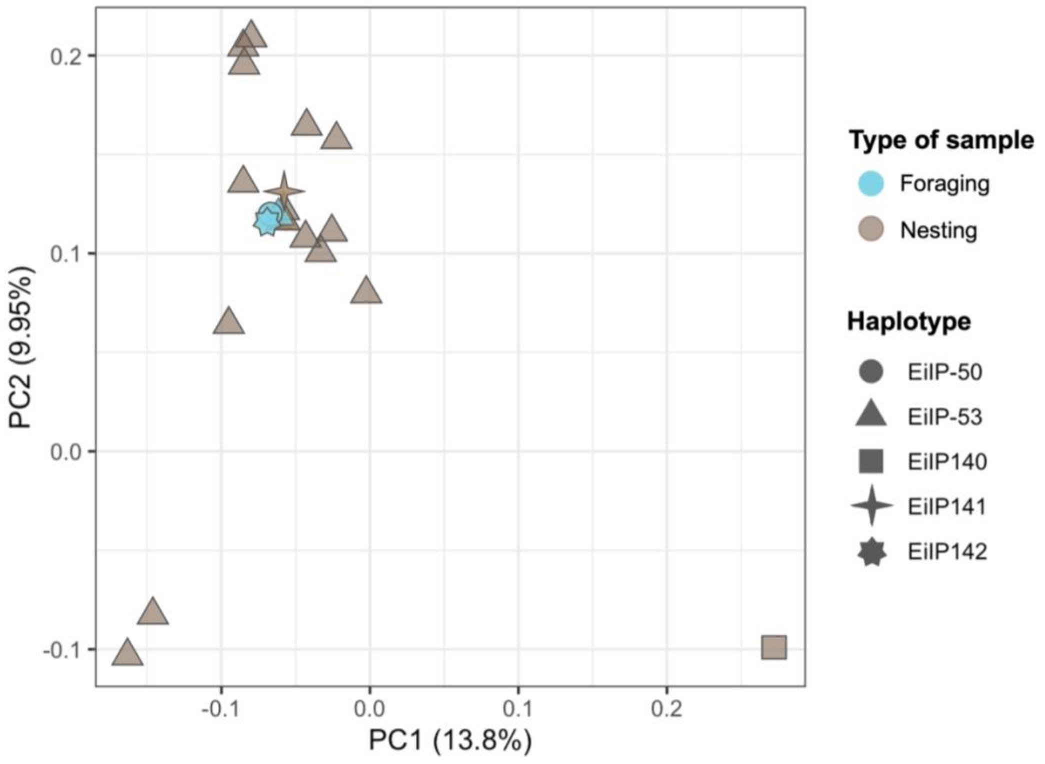


*Figure 4. Principal component analysis (PCA) of 21 unrelated hawksbill turtle genomes using filtered SNPs.*

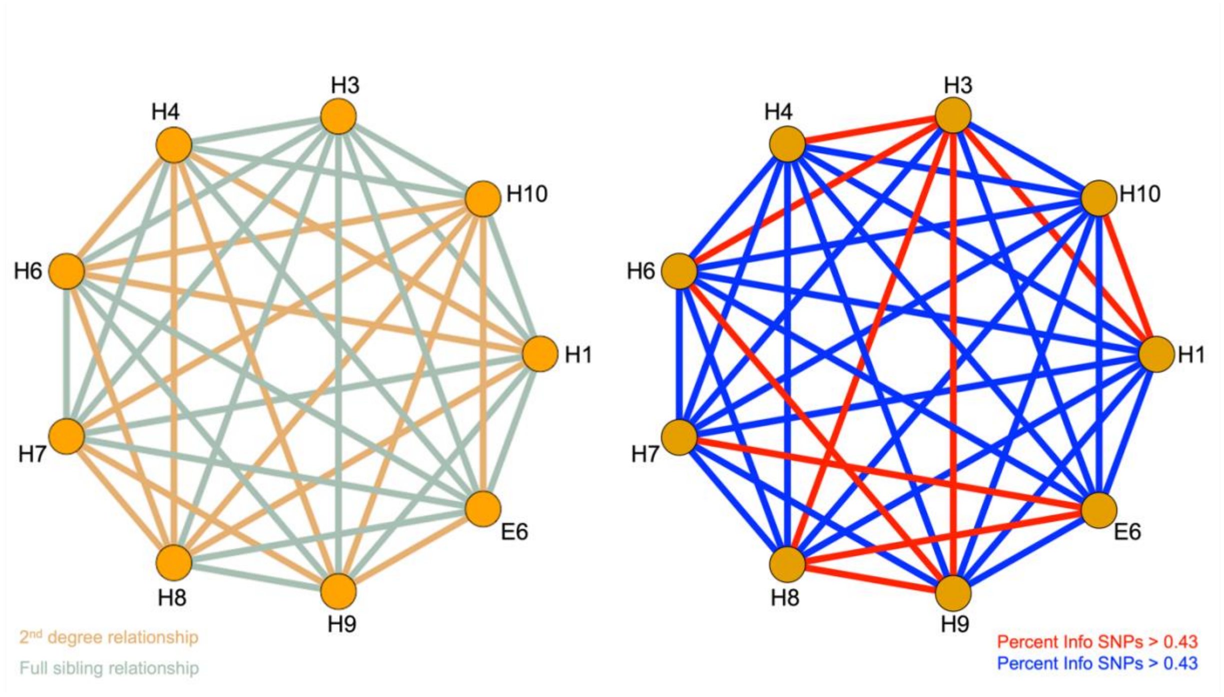


*Figure 5. Network analysis of relationships using IBS results within the nest with half-siblings.*

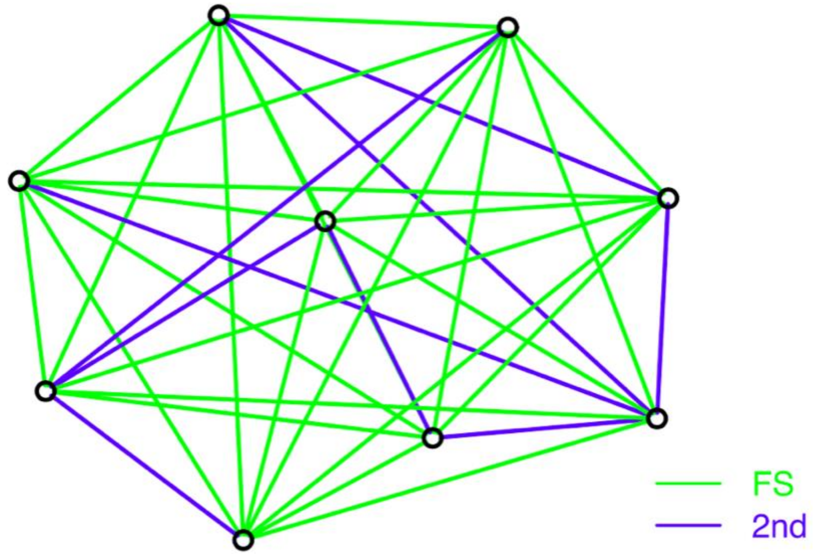


*Figure 6. Pairwise relationships among the siblings using identity by descent (IBD). FS and 2nd representing full siblings and 2nd degree relationship respectively.*
